# Evolution of *Salmonella* Chromosomes and Its Influence on Gene Expression and Chromosomal Conformation

**DOI:** 10.1101/2024.12.10.627768

**Authors:** Yueming Hu, Zewei Chen, Akosiererem S. Sokaribo, Ziyi Zhao, Xin Cao, Junya Zhang, Shuhong Liu, Runhong Chen, Yuping Deng, Hongxia Bao, Xinjie Hui, Ming-an Sun, Guo-Qiang Zhu, Shu-Lin Liu, Aaron P. White, Yejun Wang

## Abstract

*Salmonella* is one of the most important bacterial pathogens in the world, causing an estimated 120 million infections each year. There is considerable diversity, with ∼2,600 serovars, but it remains unclear how the genomes have evolved as the species and subspecies of *Salmonella* differentiate. Here, we have reconstructed the ancient orthologous chromosomes of each major *Salmonella* lineage and traced their evolutionary process. In total, 911 rearrangement events were identified, with 64% of events occurring in a locus-specific way. Using RNA sequencing and multi-strain association analysis, we demonstrate that genetic rearrangements have a significant effect on gene expression across *Salmonella* lineages. Moreover, we perform chromosome conformation capture (3C) sequencing analysis, which demonstrates large variations for the organization of *ter* chromosomal interaction domains among *Salmonella* lineages. In conclusion, our work delineates the evolutionary trajectory of *Salmonella* chromosomes, and demonstrates the influence of rearrangements on gene expression profiles and chromosomal conformation.

**Importance:** This study reconstructed the ancient orthologous chromosomes of the *Salmonella* genus, species and well-recognized subspecies, by which the trajectory of gene flow caused by genetic rearrangements was delineated chronologically. The rearrangements show apparent ‘hotspot’ distribution property and are enriched with genes important for bacterial fitness. Correlation analysis further disclosed the general influence of the rearrangements on gene expression and the organization and conformation of chromosome interaction domains. The results provide new insights on the evolution of *Salmonella* chromosomes, especially the genetic rearrangements and their epigenetic consequences.

## Introduction

*Salmonella* diverged from *Escherichia* 100 million years ago (1, 2), and then differentiated to species *S. bongori* and *S. enterica*. *S. enterica* have further diverged into 7 subspecies (3), and more *S. enterica* subspecies or subgroups were recognized most recently (4, 5). The *Salmonella* genus contains ∼2,600 distinct lineages, most of which belong to *S. enterica subspecies enterica*, with strains that can infect both cold– and warm-blood animals (6).

The phenotypic evolution and fitness of various *Salmonella* clades can be inferred from the reconstructed architecture of the ancient genomes and their evolutionary trajectories. Recently, an archaeogenomic study disclosed the genomes of eight ancient *Salmonella* strains isolated from the skeleton of humans living 6,500 years ago, which were highly similar to that of *S*. Paratyphi C (7). Archaeogenomic studies have also disclosed the ancient genomes of *Mycobacteria tuberculosis* (8), *Helicobacter pylori* (9), and other bacteria (10). However, the studies rely on the skeleton or fossil of ancient humans or other multi-cellular organisms only several thousand years ago, but the evolutionary history of bacteria is much longer. As a supplement, endeavors have been attempted to infer the more ancient genomes from the extant organisms and relevant phylogenetic information, including systematic comparison of genomes among the corresponding organisms (11–14). Despite comparative genomic studies that have been performed extensively with *Salmonella* (15, 16), there remains a lack of understanding about *Salmonella* gene flow as it pertains to the ancient evolutionary history that is unrevealed yet.

Although it is believed that genetic rearrangement can negatively influence the fitness of bacterial strains (17), large plasticity has been reported in *Salmonella* (18, 19). For example, the chromosomes of individual strains of *S. enterica subsp. diarizonae* can differ by more than 200kb in length (20). An up to 16**°** replichore imbalance of *S*. Typhimurium LT2 chromosome does not cause significant difference in growth rates and competition indices (21). Most recently, Waters *et al.,* detected genome variations in *S*. Typhi long-term culture by long-read sequencing, and found the impact of the variations on gene expression (22). These strains under investigation, however, are often phylogenetically close, and therefore the rearrangements could be transient. It is unclear whether the orthologous genomes of *Salmonella* strains belonging to different species or subspecies also show large-scale rearrangements. It is either unknown whether the rearrangements influence chromosomal conformation and gene expression.

In this study, we reconstructed the ancient orthologous genome architecture of *Salmonella* genus, species and major phylogenetic branches, based on the complete genome sequences of representative bacterial strains. Gene flow between ancestor genomes was delineated dynamically along the evolutionary history of *Salmonella* lineages. The influence of major genetic rearrangements, mainly large-scale insertions/deletions, was also observed, on the gene expression of the background genomes. Moreover, we analyzed the chromosomal topology and intra-chromosome interactions using 3C sequencing, which was developed to resolve the genome structure and inter/intra-genome interactions within bacteria and other microorganisms (23–25), but has not been applied to *Salmonella* yet.

## Results

### Reconstructing the ancient orthologous genome of *Salmonella*

To observe genome evolution, we reconstructed the ancient orthologous chromosome (AOC) of the *Salmonella* genus. Two strategies were used to build the genus AOC with the similar principles but different size of representative strains for *S. enterica subsp. enterica*. The first strategy was purely based on manual comparison and annotation, and fewer representative strains were included. Besides the 2-3 representative strains with full or nearly full sequenced genomes for each of the typically known main phylogenetic branches of non-*enterica*-subspecies *Salmonella*, only nine strains of *S. enterica subsp. enterica* were included, for which the full genomes were sequenced in earlier days and better annotated. The non-*enterica*-subspecies branches referred to *S. bongori* and six well-known subspecies of *S. enterica*, including *arizonae*, *houtenae*, *vii*, *diarizhonae*, *salamae* and *indica* (3). Except for *S. bongori*, *arizonae* and *diarizonae*, for which full genomes were available in the genome databases and selected randomly, we sequenced and assembled the full genomes for the other representative strains for the other branches (Dataset S1). The publicly available nearly-full genomes for *houtenae*, *vii*, *salamae* and *indica*, one for each of the branches, were also included for the analysis (Dataset S1). The genomes of these 26 representative strains were compared, generating a set of core genes (Dataset S2), based on which a core genomic tree was built (Fig. S1A). The tree is robust and generally consistent with previous phylogenetic results based on housekeeping genes (3). Using this tree as a guide, we manually constructed the AOC of *Salmonella* with a recursive Backbone-Patching (BP) approach and Maximum Homologous Block (MHB) algorithm newly proposed in this study (Materials and Methods).

The second strategy was based on BactAG, a semi-automatic software tool newly developed to implement the approach and algorithm constructing the AOC of a specific bacterial group. With BactAG, we re-built the genus AOC of *Salmonella*, with an overall bottom-up scheme (Materials and Methods). The AOCs of each *Salmonella* subspecies, the major ancient phylogenetic nodes between *S. enterica* subspecies and species disclosed from the core genome tree of the representative strains, and *S. bongori*, were resolved without patching by outgroups. Each AOC was built on the nearest descent orthologous chromosomes (or the extant chromosomes for the last level). The genus AOC was further patched by the genome of *E. coli* strain MG1655 and compared to the manually annotated genus AOC. With this strategy, full genomes of 85 strains representing 59 serovars of *S. enterica subsp. enterica* were included for the subspecies tree reconstruction and the subspecies AOC analysis (Dataset S1 and Fig. S1B), and therefore, a total of 102 representative *Salmonella* strains were used for construction of the genus AOC.

The genus AOCs constructed by the two strategies contained the same set of genes with the same positions and orders, suggesting the representability of the 26 strains included for manual analysis and the overall accuracy of the AOCs (Fig. 1A; http://61.160.194.165:3080/ESG/datasets/AG). For convenience, the manually annotated genus AOC was used for subsequent analysis, and the nucleotide sequence for each orthologous block of the AOC was represented by the corresponding sequence retrieved from a randomly selected strain under comparison that contains the orthologous region. Consistent with the GC skew (26) shifting points, the replication origin (*oriC*) and terminus (*terC*) separate the genus AOC into two halves with nearly identical size (Fig. 1A).

**Fig. 1.**
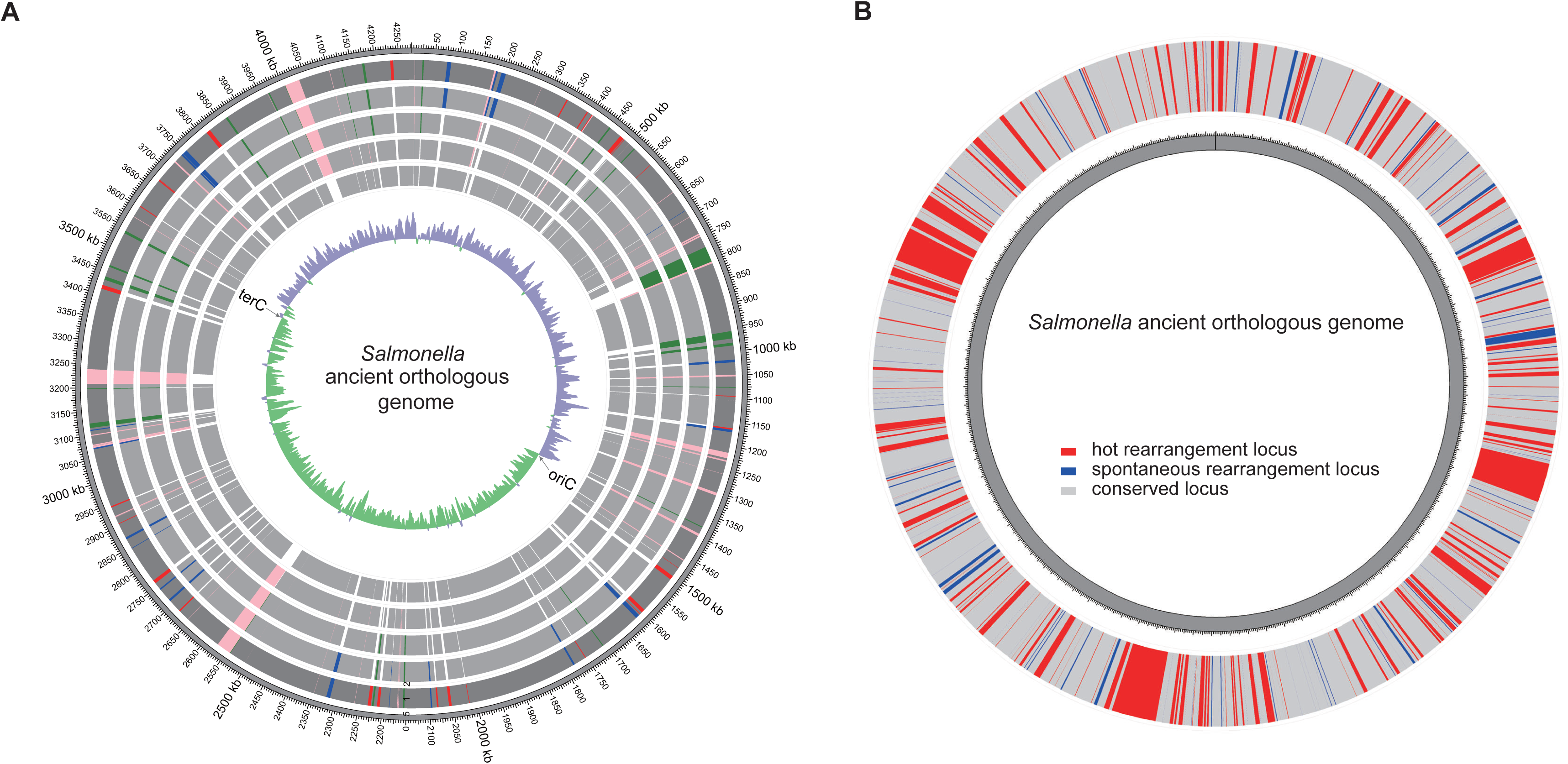
The ancient orthologous chromosome of the *Salmonella* genus and distribution of genetic rearrangements. **(A)** The ancient orthologous genome frame of *Salmonella* genus. The genome frame was inferred manually, with the sequence of each orthologous block represented by that of any randomly selected strain under comparison. The 2^nd^ to 6^th^ circles from inside to outside represent the resolving process of the ancient orthologous genome of *Salmonella*: the primary backbone fragment (in grey) of *Salmonella* ancient orthologous chromosome inferred by the genome comparison between *S*. typhimurium LT2 and *S*. *bongori* N268-08 (the 2^nd^ circle), the updated blocks of *Salmonella* ancient orthologous chromosome inferred by the genome comparison between LT2 and other representative *S*. *bongori* strains (in pink, the 3^rd^ circle), between the representative strains of various subspecies from *S. enterica* and *S*. *bongori* strains (in green, the 4^th^ circle), between the representative strains of *E. coli* MG1655 and *S*. typhimurium LT2 (in blue, the 5^th^ circle) and between MG1655 and *S*. *bongori* N268-08 (in red, the 6^th^ circle). The 6^th^ circle also represents the final ancient orthologous chromosome of *Salmonella* genus indicated with the primary backbone in dark grey and the updated blocks in the corresponding colors described above. The innermost circle is the GC-skew curve of the ancient orthologous genome of *Salmonella* genus. The chromosome replication origin and termination sites were predicted by the GC-skew results and alignment against LT2 chromosome, and noted as ‘*oriC*’ and ‘*terC*’ respectively. **(B)** Distribution of rearrangement loci in the ancient orthologous chromosome of *Salmonella* genus. The ‘hotspots’ and ‘spontaneous loci’ were shown in red and blue lines respectively.

### The evolutionary trajectory of *Salmonella* chromosomes

BactAG was based to construct the AOCs corresponding to the major *Salmonella* phylogenetic branch (http://61.160.194.165:3080/ESG/). By comparison between neighbor AOCs, we identified 911 rearrangement events of >1kb from the evolutionary trajectory, which were unevenly distributed along the chromosomes (Fig. 1B). These loci were further classified into “spontaneous”, where only one rearrangement event was identified along the evolutionary trajectory, and “hot spots” where rearrangement events happened repeatedly. There were more hot spots (64%) than spontaneous rearrangement loci (36%) (Fig. 1B; Fig. S2A), indicating that *Salmonella* chromosomal rearrangements may have happened in a locus-dependent way (27, 28). In total, 253 known functional entities were annotated from the genes encoded in these rearrangement loci (Fig. 2A), many of which are associated with virulence (88, 35%), fimbriae biosynthesis (47, 19%) and iron uptake (13, 5%) (Fig. S2B).

**Fig. 2.**
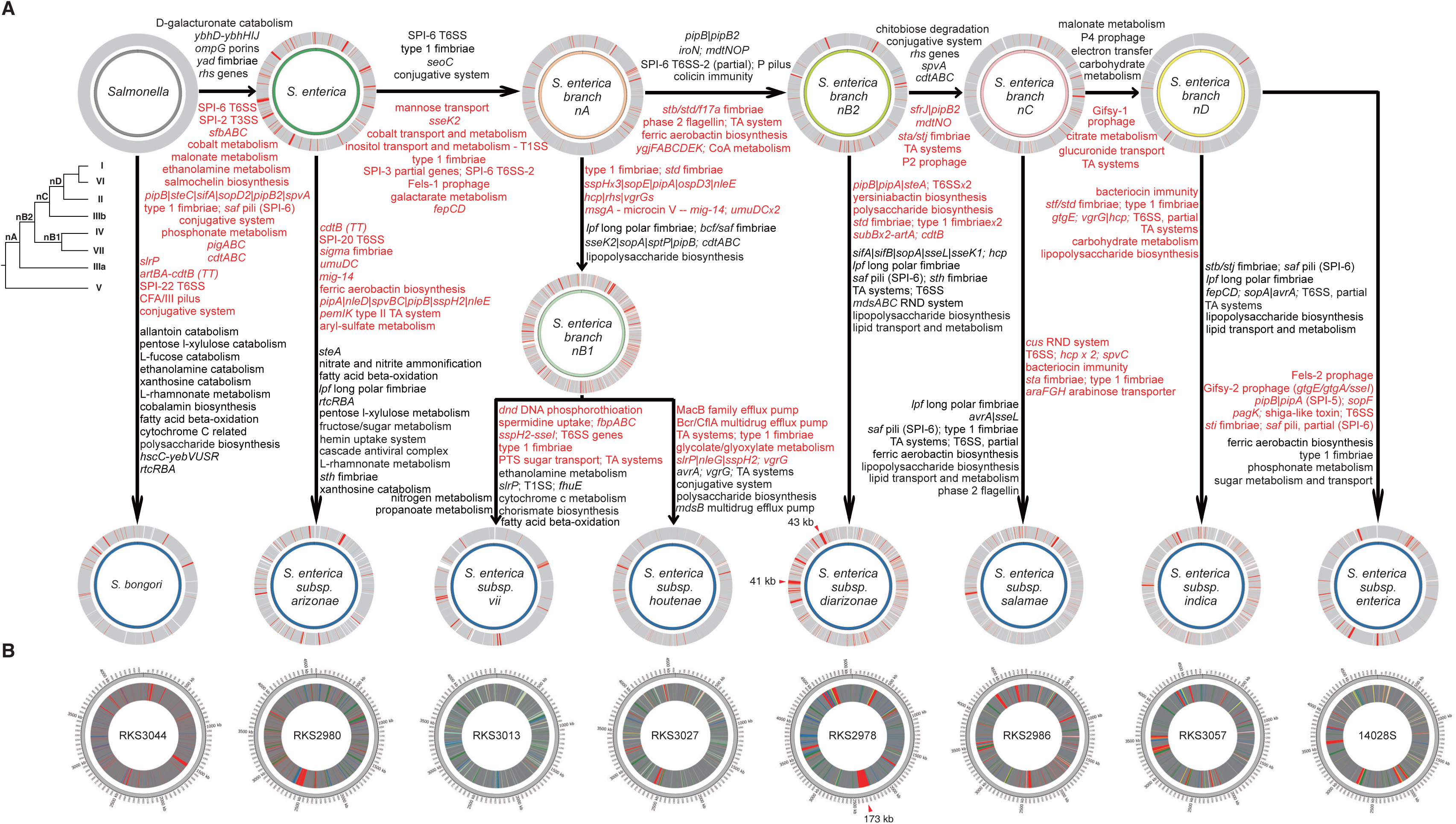
The evolutionary trajectory of *Salmonella* ancient orthologous chromosomes. **(A)** Gene flow along the evolutionary trajectory of *Salmonella* chromosomes. Each circle represents the ancient orthologous chromosomes of different *Salmonella* phylogenetic nodes, and the general trajectory is shown with directed lines according to the phylogenomic tree shown on the left. At the end of each branch, the genome circles show regions conserved with the genome of the parent node in grey, gained in red or lost in white. Two blocks >40kb that were gained in *S. enterica subsp. diarizonae* (IIIb) are indicated with arrows. Beside each arrow connecting the different parts of the phylogenetic tree, the representative operons or pathways acquired from the parent nodes are listed in red, whereas those lost are listed in black. **(B)** One-dimensional representation of the chromosomes of representative strains with phylogenetic blocks. The blocks of a certain phylogenetic origin were shown in the color consistent with that of the inner circle for the corresponding phylogenetic node. The arrow in the RKS2978 chromosome represents a strain-specific block of 173kb.

Evolutionary routes can be traced for individual genes according to the gene flow chart analyzed from the reconstructed ancient orthologous genomes (Fig. 2A). For example, the gene flow chart analysis suggests that, the *lpfABCDE* (*lpf*) operon encoding long polar fimbriae distributed in only a few *Salmonella* strains (29) was anciently present but lost subsequently in the ancestors of different branches of *S. enterica* (Fig. S3). Like *lpf*, the evolutionary routes of *cdtABC* (30) (Fig S4), the operon encoding typhoid toxin (31) (Fig. S5) and 21 other genes or operons were analyzed according to the evolutionary trajectory of ancient orthologous genomes (Dataset S3).

By comparison to the ancient orthologous genomes of major nodes, we fragmentally represented the chromosomes of individual strains with evolutionary origin, following a one-dimension genome representation (1DGR) annotation pipeline (Materials and Methods; Fig. 2B). In ancient orthologous genomes, only two DNA insertions were >40kb in length, and both were present in the ancient orthologous genome of *S. enterica subsp. diarizonae* (41 and 43kb; Fig. 2A). In strains at the end of the branchpoints, however, the rearrangements happen more extensively with larger size of rearranged fragments, as exemplified by the 173-kb insertion in *S. enterica subsp. diarizonae* strain RKS2978 (Fig. 2B). Therefore, the previously reported large plasticity of *Salmonella* chromosomes could be mainly constrained to the recently diversified phylogenies (18, 19, 32), which would eventually be purified after long-term selection and fixation to maintain a conserved form.

### Influence of chromosomal rearrangement on *Salmonella* gene expression

To observe the influence of the identified genomic rearrangements on gene expression, we performed RNA-seq analysis on eight representative *Salmonella* strains, one for each species and subspecies, at both exponential and stationary phases of growth (Dataset S4). When all differentially expressed genes were considered, strain clustering was so strong that the two growth phases could not be distinguished at all (Fig. 3A). However, when we focused solely on differentially expressed genes that were part of the core gene set, there was an obvious split between the exponential and stationary phases of growth (Fig. 3B). This clustering pattern indicated that the unique genes within a strain were driving the genetic divergence. The number of differentially expressed core genes is no longer smaller between growth stages than between strains (Fig. S6). However, there remain hundreds of conserved genes with significant expression difference between strains at the same growth phase, suggesting potential effects of genetic rearrangements.

**Fig. 3.**
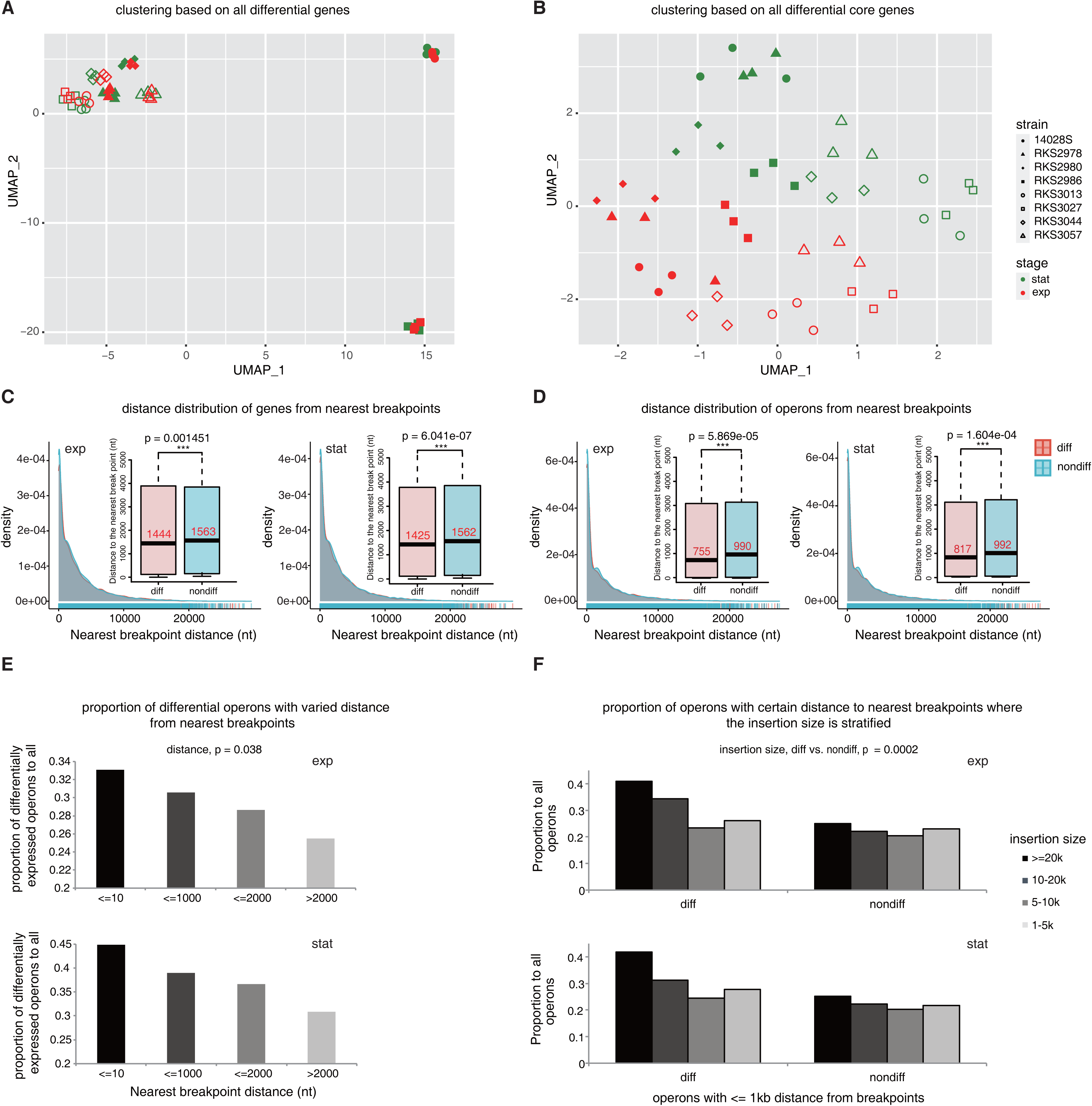
Influence of genetic rearrangements on *Salmonella* global gene expression profiles. Clustering of RNA-seq samples based on all differentially expressed genes **(A)** or just genes that are part of the *Salmonella* core genome **(B)**. UMAP was used for reducing the dimensions and visualization (Materials and Methods). **(C)** – **(D)** The distances away from the nearest upstream 1DGR breakpoints in the ancestral genomes for each strain were calculated at each growth phase for differentially (‘D’) and non-differentially (‘ND’) expressed genes **(C)** or operons **(D)**. The *p*-values of Mann–Whitney U tests are shown and statistical significance is indicated with asterisks. **(E)** The proportion of differentially expressed operons at each growth phase was calculated based on the distances away from the nearest upstream 1DGR breakpoints. One-way ANCOVA was performed with the growth stage as a covariate, and the *p*-value for equal distribution of proportion among distance strata was indicated. **(F)** The proportion of differentially expressed or non-differentially expressed operons less than 1000bp away from the nearest breakpoints was calculated based on the insertion block size at the breakpoints. Two-way ANCOVA was performed with the growth phase (i.e., exp – exponential; stat – stationary) as a covariate; the *p*-value for the trend in proportion distribution based on the insertion block size is listed.

We traced the evolutionary origins of the differentially expressed genes, and found that most of them are conserved, with the origin of genus ancestor (Dataset S5). All genes originating from the genus ancestor were further retrieved, and classified into differentially and non-differentially expressed groups. We compared the distances from these groups of genes to the nearest breakpoints where rearrangements had occurred (Materials and Methods). On average, the differentially expressed operons and genes were physically closer to rearrangement spots than were the non-differentially expressed operons and genes (Fig. 3C-D). At rearrangement positions where large insertions >1kb had occurred, we found that a higher proportion of operons were differentially expressed, for both growth phases tested (Fig. 3E). We also compared the size of the insertions and found that larger insertions had a higher proportion of differentially expressed genes close to them and the proportion dropped based on the size of the insertion (Fig. 3F). Taken together, the results indicated that rearrangements had a strong effect on expression of neighboring genes within the backbone genome, and the influence was dependent on the distance from the breakpoints and the size of the rearrangements.

### Conservation of chromosomal overall conformation in *Salmonella*

We also observed the influence of rearrangements on the conformation of *Salmonella* chromosomes. 3C experiments were performed on exponentially grown cells from representative strains of *S. bongori* (RKS3044), and *S. enterica subsp. arizonae* (RKS2980), *houtenae* (RKS3027) and *indica* (RKS3057). The contact maps showed a high correlation among biological replicates (Fig. S7), with all displaying a single strong diagonal that demonstrates the enrichment of contacts between neighboring loci as in *E. coli* (25) (Fig. 4A). Scalograms demonstrate a pattern of *ter*-centered constraint and *ori*-centered looseness of DNA-DNA contact organization for all the *Salmonella* chromosomes (Fig. 4B). Similar to *E. coli* (25, 33, 34), six conserved major intervals can be identified from the chromosomes of all the *Salmonella* strains, as suggested by Directional Index analysis (Fig. 4C). The boundaries of most *Salmonella* major intervals show homology to those of *E. coli* MG1655, especially for *ori* and the adjacent intervals (Fig. S8A-B). Consensus sequence motifs can be identified from these boundaries (Fig. S9 and Dataset S6).

**Fig. 4.**
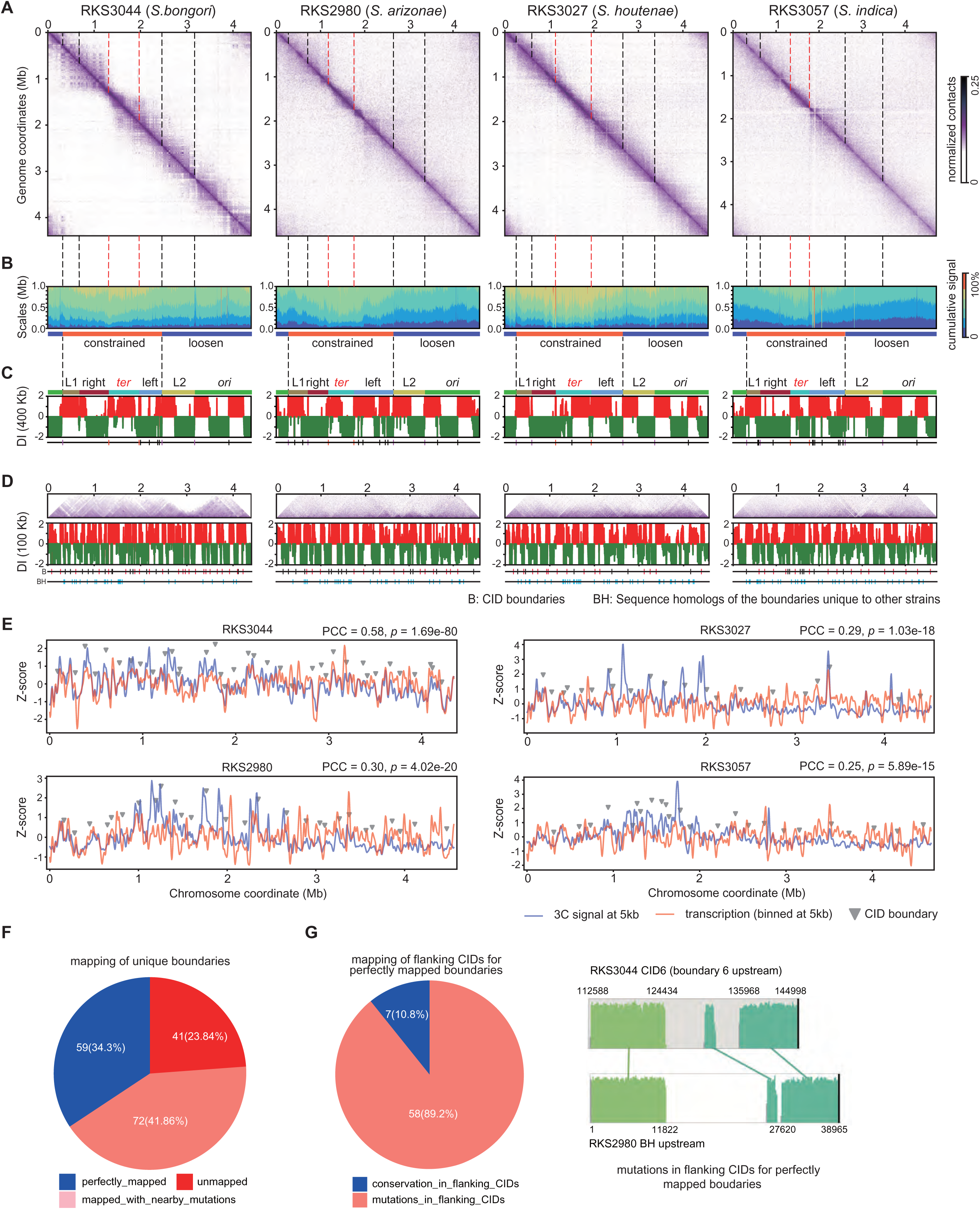
Conformation of *Salmonella* chromosomes. **(A)** 3C contact maps of RKS3044, RKS2980, RKS3027 and RKS3057 chromosomes. The bin size was set as 5kb. The reference chromosome sequence for each *Salmonella* strain was adjusted to match the chromosomal annotation start site and strand referred to from *E. coli* strain MG1655. **(B)** Scalogram representation of chromosomal contact maps of the representative *Salmonella* strains. **(C)** 400-kb Directionality index (DI) analysis of chromosomal contact map of the representative *Salmonella* strains. The boundaries of major intervals analyzed according to the 400-kb DI results were indicated below the DI diagrams. **(D)** The organization and boundaries of *Salmonella* CIDs analyzed according to 100-kb DI results. B: boundaries of CIDs, which were represented with red or black vertical lines if they are or not conserved in at least another *Salmonella* or *E. coli* strain respectively. BH: distribution of sequence homologs of CID boundaries of other *Salmonella* strains or MG1655 but absent in the observed strain. **(E)** Correlation between gene transcription levels and short-range contact frequencies. The RNA-seq read count and DNA-DNA contact pairs were combined within 50-kb bins along each chromosome and normalized. The CID boundaries for each strain was shown with arrows. Pearson correlation coefficients (PCCs) and *p*-values were indicated. **(F)** Sequence conservation of strain-unique CID boundaries in other *Salmonella* strains. **(G)** Sequence conservation of CIDs flanking the strain-unique boundaries that could find sequence homologs in other *Salmonella* strains. An example was shown to illustrate the extensive rearrangements in the flanking CIDs.

### Evolution of the *ter* major interval in *Salmonella* chromosomes

In contrast, the *ter* interaction interval varied in size (Fig. 4C) and had non-conserved boundaries (Fig. S8A-B). Comparative genomics identified a >500-kb inversion spanning the *ter* site between MG1655 and all the four *Salmonella* strains analyzed (Fig. S8C-D). Besides MG1655, the inversion was also detected between other strains covering various *E. coli* lineages (35, 36) and *Salmonella* strains (Dataset S7). Notably, the boundaries of the RKS3057 *ter* interval coincide with those of MG1655 with cross-homology, i.e., the upper *ter* boundary in RKS3057 being reverse-complementarily homologous to the lower boundary in MG1655, vice versa (Fig. S8B-C). The boundaries in MG1655 and RKS3057 are located at the ends of the inversion (Fig. S8C-D). The lower *ter* boundary in RKS2980 also shows cross-homology to that in MG1655 (Fig. S8C-D). The interpretation of this is that the large inversion between *E. coli* and *Salmonella* potentially happened anciently before the diversification of *Salmonella* species, was *ter*-centered, and did not change the *ter* interaction interval or the boundaries when the event happened. However, the boundaries could have since varied in subsequent phylogenies. The variation does not appear to be caused by rearrangements of boundary sequences, which are conserved in all the examined strains (Fig. S8C-D), but might be related to significant rearrangements in the flanking regions. There is a large rearrangement-hotspot block within the *ter* interval of *Salmonella* ancient chromosomes (Fig. S10).

### Diversified organization and conserved boundary sequence patterns of chromosome interaction domains in *Salmonella*

The number of chromosome interaction domains (CIDs) varied among strains RKS3044 (35), RKS2980 (28), RKS3027 (21) and RKS3057 (24), and the sizes of CIDs varied from 15kb to 775kb with a median length of 150kb (Fig. 4D). 62% (23/37) for RKS3044, 71% (15/21) for RKS3027, 67% (16/24) for RKS3057 and 46% (13/28) for RKS2980 of the CID boundaries were conversed in at least one other *Salmonella* strain or *E. coli* MG1655 (Fig. 4D; Fig. S11A). The boundary sequences of some unconserved major intervals could form non-major-interval CID boundaries in other chromosomes. For example, the lower *ter* boundaries of RKS3057/RKS2980 are homologous to the boundary of a CID in RKS3044 or RKS3027 (Fig. S11A, arrow). Consensus motifs were identified from the conserved CID boundaries (Fig. S12A-B) or the pooled conserved boundaries of CIDs and major intervals (Fig. S12C-D). One motif frequently detected from the boundaries of both major intervals (reverse complementarily) and CIDs, ‘YGGCGCTGGM’ (Dataset S6), is homologous to the binding motif of RpoN (37) (https://meme-suite.org/meme). Consistent with the observations in other bacteria (25, 38, 39), for all the four *Salmonella* strains, the abundance of transcribed RNAs showed significant correlation with the short-range (within 5kb) DNA-DNA contact frequency along the genome (Fig. 4E). The CID boundaries were often located nearby the peaks of transcription read count (Fig. 4E).

### Influence of genetic rearrangements on *Salmonella* chromosome interaction domains

Each *Salmonella* chromosome also shows unique CID boundaries, which are particularly enriched in *ter* or the adjacent major intervals (Fig. S11A). In RKS3044, 12/14 of the unique CID boundaries are located in *ter*-centered constrained region (Fig. S11A; *p* = 0.003; χ^2^ test). However, most of the strain-unique CID boundaries can find their sequence homologs in other strains (BH; Fig. 4D; Fig. S11B taking RKS3044 as an example). In total, only 41 of 172 (23.8%) of strain-unique boundaries did not have sequence homologs in the other strains (Fig. 4F). Homologs can be found for more unique boundaries but with striking nearby rearrangements, i.e., >1kb rearrangements within 10kb flanking regions (72/172, 41.9%; Figure 4F). For 89% (58/65) of the non-boundary sequence fragments that are perfectly mapped by the unique boundaries of other strains, the flanking CIDs show large (>10kb) fragmental rearrangements (Fig. 4G). The results suggest that, besides local sequence features, nearby rearrangements can also influence the formation of CID boundaries. As an example, the boundaries of a CID where the *Salmonella* Pathogenicity Island 2 (SPI-2) Type III Secretion System (T3SS) (40–42) is located demonstrate how rearrangements are associated with changes of local chromosomal conformation (Fig. S13A-C).

### Long-range intra-chromosomal interaction profiles of *Salmonella* strains

The rearrangements also influenced long-range DNA-DNA interactions. Among the long-range interactions identified by 3C analysis, which were defined with >50-kb spacing distance (Materials and Methods), the top-ranked loci with the largest number of interacting pairs were consistent among *Salmonella* strains for the function of encoding genes, such as those encoding SPI-1 and SPI-2 T3SSs and effectors, other virulence factors, fimbriae or curli. The individual loci repeatedly present in different strains with widest interactions were retrieved, including the SPI-1/2 T3SSs and others (Fig. S14A-D). The loci encoding SPI-2 T3SS and related effectors are missed in RKS3044, and the long-range interactions involved in these equivalent loci are much sparser and more concentrated than in *S. enterica* strains (Fig. S14A vs. S14B-D). The long-range interaction profiles involving the SPI-1 T3SS or effector-encoding loci also showed striking differences, especially between *S. bongori* (RKS3044) and the *S. enterica* strains, partially associated with genetic rearrangements (Fig. S14A-D). For example, the RKS3044-specific *nleG* locus shows wide and long-range interaction connecting the loci of *rtsAB*/T1SSs and *csg* directly or indirectly (Fig. S14A). Another RKS3044-specific locus, *slrP-artBA-cdtB*, connects *csg* and the SPI-2 T3SS equivalent locus (Fig. S14A).

## Discussion

There are contradictory observations for the evolution of *Salmonella* chromosomes. Phylogenetically distant strains, e.g., *S*. typhimurium and *S. bongori* strains, show large conservation of gene composition and order in chromosomes (43, 44). In contrast, extensive and large-scale rearrangements have been found among closely related strains exemplified by *S*. typhi and others (19, 32, 45). In this study, by tracing and comparing the ancient orthologous chromosomes, we confirmed that the ancient orthologous *Salmonella* chromosomes before the divergence of subspecies show much less extensive recombination than the extant strains compared to the ancestors (Fig. 2). Besides the scale constraint of local rearrangements, the long-term conservation of *Salmonella* chromosomes could be further maintained by the rearrangement pattern of hotspots (Fig. 1B), which was also observed previously (27, 28). Rearrangements could happen with scale constraint and repeatedly in the preferred loci with a limited total number, leading to the conservation of *Salmonella* chromosomes during the long-term evolutionary process. It should be noted that we analyzed the ancient orthologous chromosomes rather than ancient chromosomes. Ancient chromosomes could temporarily contain more horizontally transferred DNA fragments, which would be lost eventually for fitting. A recent study confirmed the presence of large rearrangements within *S*. Typhi population under culture, and the rearrangements showed apparent influence on both gene expression and bacterial fitness (22). Therefore, the larger-scale rearrangements observed in the chromosomes of extant strains could reflect the temporal evolution rather than real chromosomal plasticity which should be defined with the premise of fitness.

The underlying mechanisms have not been investigated by which *Salmonella* chromosomes keep the large conservation during the long period of evolution. Large-scale rearrangements could cause deleterious influence on gene expression profile or chromosomal conformation, and therefore are purified during the long-term evolution (17, 18, 46, 47). To validate the hypothesis, gene expression profile and chromosomal conformation should be observed for multiple *Salmonella* strains of various phylogenetic origins. In this study, we made the first attempt to perform both the gene expression profiling studies and 3C experiments for the representative strains of multiple *Salmonella* species and subspecies, generating important resources for the research community.

The *Salmonella* strains with different species or subspecies origins showed exceptionally large difference in gene expression (Fig. S6A). Unsupervised clustering of bacterial samples featured by all the differentially expressed genes demonstrated larger influence of genetics than growing phases on general gene expression profile (Fig. 3A). When only the genetically conserved genes were investigated, there was totally different observation, i.e., larger influence on expression of conserved genes exerted by phase shifting than genetic variation (Fig. 3B). In other words, the general expression of conserved genes within *Salmonella* background chromosome turns to be stable among genetically different strains. However, rearrangements still show influence on the expression of conserved genes, and the influence depends on the distance of rearrangements away from the targeted genes and the scale of rearrangements (Fig. 3C-F). The expression of conserved genes within *Salmonella* background chromosome, which are potentially required for maintenance of bacterial basic life processes, could be an important constraint factor, whose changes may cause bacterial fitting problems and therefore the scale and frequency of rearrangements are constrained.

The conformation could be another important constraint factor to maintain the long-term conservation of *Salmonella* chromosomes. Diverged before ∼100 million years, the conformation of chromosomes between *Salmonella* and *E. coli* or between *Salmonella* branches appears super-conserved, as reflected by the conservation of boundaries and boundary sequences for most major interaction intervals (Fig. 4A-C). Despite a ∼600 kb inversion spanning *ter* between the ancestor of *Salmonella* and *E. coli*, organization or the boundaries of *ter* intervals was not influenced (Fig. S8, MG1655 vs. RKS3057). We cannot even exclude the possibility that the un-conservation of partial major-interval boundaries in some strains may represent the short-term evolutionary outcome of individual strains rather than the stable status after long-term adaptation. It means that the conservation of general conformation of *Salmonella* chromosomes is possibly beyond what we have observed. Different from the major intervals, the number and organization of CIDs within each major interval show large variance from *Salmonella* strains to strains of different evolutionary origins (Fig. 4D). We noticed that the boundaries for many CIDs are not conserved but the sequences seldom mutate extensively among strains, and yet the boundary extension or pre-stop of the CIDs is associated with apparent rearrangements within boundary-flanking regions frequently (Fig. 4E-F). Therefore, rearrangements have important influence on the formation and organization of CIDs. We also found that the genes frequently involved in rearrangements in *Salmonella* chromosomes, e.g., genes encoding SPI-1 or SPI-2 T3SSs and their effectors, virulence genes, iron transporter-encoding genes and fimbriae genes, also represent the hub loci of wide interactions with other far-spanned chromosomal loci, which could lay the important foundation for the CID organization variations (Fig. S14). Taking together, the conservation of general conformation of chromosomes could be another important factor maintaining the chromosomal conservation and constraining the total scale of genetic rearrangements, while the influence of rearrangements on local chromosomal conformation could be adjusted by the frequent rearrangements happening in rearrangement hotspots during the long-term adaptation process.

In summary, the study systematically investigated the general evolutionary property of *Salmonella* chromosomes. However, there are more unknowns requires more well-designed experiments to clarify. Associations were observed between genetic rearrangements and the expression of conserved genes, and the relationship was also evidenced by other studies (48), but the definitive mechanisms connecting the two molecular phenotypes remain unclear. Rearrangements could be the major factors themselves (22), but the influence could also be caused by microevolutions such as SNPs or short indels. Expression changes of conserved genes could be caused by direct influence on their *cis* elements, indirect influence by acting on their *trans* factors, or epigenetic influence through changing the chromosomal conformation. In this study, we also found the conservation of general conformation and the boundary sequences of *Salmonella* chromosomes; however, we remain unclear whether the sequences play a determining role and how they function. Consistent with other studies (38, 39), we also found significant correlation in different *Salmonella* lineages between gene expression and CID organization along chromosomes (Fig. 4E). However, it is unclear whether there is mediating effect for gene expression between genetic rearrangements and chromosomal conformation, for chromosomal conformation between genetic rearrangements and gene expression, or both. There are also other interesting questions that are yet to clarify. For instances, have the chromosomes of some *Salmonella* strains (e.g., *S*. typhi), which show large-scale inversions and translocations compared to other *Salmonella* chromosomes (19, 32, 45), undergone general conformation changes? If so, do the chromosomal conformation changes lead to fitting problems? Compared to NTS, do the more rapid microevolutions in the genomes of iNTS strains influence chromosomal conformation? Furthermore, is the chromosomal conformation conserved or different between the bacterial strains with the same genetic background but with different states and gene expression profile, for example, between the same strains grown to different phases (49), or between the planktonic and aggregated states (50)? To solve the problems could facilitate understanding the evolution of *Salmonella* chromosomes more clearly.

## Conclusions

Taken together, in this study, we have delineated the evolutionary trajectory of *Salmonella* AOCs, and have demonstrated that genetic rearrangements can have significant effects on global gene expression profiles, as well as chromosomal conformation.

## Materials and Methods

### Genome sequencing, assembly and annotation

The genomes of five *Salmonella* strains were sequenced in this study (Dataset S1). The strains were inoculated in Luria-Bertani (LB) agar plates and incubated at 37℃ overnight, and single colonies were inoculated in LB medium and grown at 37℃, shaking at 200rpm for 8 hours. For each strain, 4ul of the culture was transferred to 200ml fresh LB medium and grown at 37℃ and 200rpm to logarithmic phase. The cells were collected and genomic DNA was isolated following a standard phenol-chloroform extraction procedure. DNA libraries of an insert size of 350-400 bp were prepared using the TruSeq DNA PCR-Free Library Kits, and paired-end 250-bp sequencing was performed on the Hiseq 2500 platform. PacBio RS II platform was also used to sequence the genomes, with an insert size of 10 kb. P6-C5 kits were used for preparation of the PacBio libraries. Spades (hybridSpades version) was used to perform hybrid assemblies based on both the short and long reads (51). RASTtk v1.3.0 was used to annotate the gene composition and function (52).

### Core genome and phylogenomic analysis

The core genome and phylogenomic analysis was performed for both *Salmonella* genus with genomes of 26 strains representing the major *Salmonella* phylogenetic branches, and *Salmonella enterica subsp. enterica* with the representative genomes of 59 subspecies serovars and a *Salmonella enterica subsp. indica* strain RKS3057 as the outgroup (Dataset S1). The publicly available genomes not sequenced in the study were retrieved and downloaded from the NCBI Genome database with the accessions listed (Dataset S1). Core genomes were analyzed with BactCG (http://61.160.194.165:3080/ESG/tools/BactCG). Briefly, protein-encoding genes were annotated from each genome, and the derived protein sequences were aligned between each pair of strains with BLASTP (https://blast.ncbi.nlm.nih.gov). Homolog pairs were identified with similarity >70% for ≥70% coverage of the full length of each aligned protein. Best mutual alignment was used to determine the orthologous pairs. A core-genome protein family was defined only if the corresponding protein contained a unique ortholog in each strain. The protein sequences of each family belonging to the core genome were aligned among the strains using CLUSTALW with the default parameter setting (http://www.clustal.org/) (53), and the alignment results of all the core-genome protein families were concatenated for each strain according to a fixed family order, followed by phylogenomic analysis using MEGA6 (54). Both Maximum likelihood (ML) and Neighbor joining (NJ) methods were used for building the phylogenetic trees. Only the NJ tree was illustrated in the study because the NJ and ML trees appeared to be consistent between each other. The Jones-Taylor-Thornton model for amino acid substitutions was used for ML tree building. Bootstrapping tests were performed for 1,000 replicates to assess the robustness of nodes; only branches with 50 percentages of the subtrees were considered stable.

### Inference of the ancient orthologous genome frame of *Salmonella* lineages

The ancient orthologous chromosome (AOC) of *Salmonella* was inferred both manually and semi-manually. Both strategies were based on a two-step Backbone-Patching approach. In the Backbone step, the most anciently diverged clades were identified according to the phylogenomic tree with strains covering the major branches of the genus, species or subspecies to be studied, and one representative strain was selected randomly from either clade. Orthologous fragments were analyzed with Mauve v2.4.0 (55) and an iterative Maximum Homologous Block (MHB) algorithm, and combined to generate the backbone of the ancient orthologous genome. Two patching sub-steps followed. Firstly, genomes of other representative strains were aligned between the two clades, and the orthologous fragments were retrieved, which were further compared to the backbone. The sub-fragments not covered by the backbone genome were patched in manually and the backbone ancient orthologous genome was updated iteratively. Secondly, the genomes of closely-related outgroup strains or the genome of nearest ancestor were also aligned against the representative strains of either clade respectively and the orthologous fragments were extracted to further patch the ancient orthologous genome. The MHB algorithm proposed in the Backbone step was to position the orthologous blocks between the two bacterial chromosomes that were compared. There were various rearrangement events in bacterial genomes so that homologous blocks between two chromosomes were not necessarily orthologous to each other. To resolve the orthologous backbone between two chromosomes, we numbered the homologous blocks in each strain according to their within-genome location, and ordered them according to the length of each block. The positioning relationship of the longest (*l*_1_) and the second (*l*_2_) longest blocks was observed in either genome and compared. If it was consistent between the two genomes, as was most often, the third longest block pair (*l*_3_) was included and the positioning relationship between *l*_3_ and *l*_1_ / *l*_2_ was compared between the two genomes. The *l*_3_ was retained if the relationship was consistent between genomes and removed if inconsistent. The composition and positional order of the homologous backbone blocks were updated subsequently. The *l*_4_ block pair would be further included and the nearest upstream (*l*_up_) and downstream (*l*_down_) blocks were retrieved from the retained longer blocks in either genome. The homology and position-ordering relationship of *l*_4_, *l*_up_ and *l*_down_ were compared between genomes and *l*_4_ was retained if consistent and removed if inconsistent. The steps were performed iteratively till the shortest homologous block pair was analyzed.

A GO package, BactAG, was developed to facilitate semi-automatic inference of the ancient genomes (http://61.160.194.165:3080/ESG/tools/BactAG). The ancient genome architecture was analyzed with the tool for each major *Salmonella* phylogenetic node. The AOC analysis for each subspecies with BactAG was performed with a procedure similar with that used for the manual analysis of *Salmonella* genus AOC. For other nodes, a bottom-up strategy was used, that is, the AOCs of sub-nodes close to the target node were used and compared directly, instead of the full set of representative strains.

### Annotation of ancient orthologous genomes

PGAG v2020-06-10.build4646 was used to annotate the ancient orthologous genomes (56). To facilitate the comparison of different genomes, we proposed PGA, i.e., a <u>p</u>an-<u>g</u>enome based <u>a</u>nnotation pipeline, to unify the gene annotation results of ancient orthologous genomes (http://61.160.194.165:3080/ESG/tools/BactPGA). With the pipeline, the *Salmonella* pan orthologous gene set was calculated from 26 representative strains by a completely combinatorial, pairwise mutual best alignment-based strategy with a minimal similarity cutoff of 0.7 for ≥ 70% full-length coverage for each aligned protein (http://61.160.194.165:3080/ESG/tools/BactPG). Furthermore, the protein-encoding genes from the *Salmonella* ancient orthologous genomes annotated with PGAG were aligned against the pan gene set and the orthologous groups were determined, followed by annotation with the unique accessions of the corresponding orthologous clusters.

### Comparison of ancient orthologous genomes

MAUVE v2.4.0 was used to align the ancient genomes of phylogenetic neighbor nodes (55). Taking the genome of the parent node as a reference, the insertions, deletions, translocations or inversions ≥1 kb in the genome of the daughter node were recorded, and annotated for the gene composition and function according to the PGA annotation files.

### Mosaic block representation of bacterial genomes with evolutionary origins

A One-Dimension Genome Representation (1DGR) pipeline was proposed to delineate the chromosome sequences of *Salmonella* strains as mosaic blocks indicated with their evolutionary origins (http://61.160.194.165:3080/ESG/tools/Bact1DGR). Briefly, for each strain, the chromosomal sequence was aligned against all the ancient genomes traced in the study, and the sequence blocks were marked with the most ancient origins according to the orthologous hits. The blocks that did not have any orthologous hits with the ancestor genomes were noted as sequence fragments specific to the strain.

### Bacterial RNA extraction, sequencing and preprocessing

Eight representative strains, including *S. bongori* RKS3044, *S. arizonae* RKS2980, *S. houtenae* RKS3027, *S. vii* RKS3013, *S. diarizonae* RKS2978, *S. salamae* RKS2986, *S. indica* RKS3057 and *S. enterica* serovar Typhimurim 14028S, were inoculated and grown in LB medium at 37℃, shaking at 200 rpm for 8 h (exponential phase) or 18 h (stationary phase); three biological replicates were prepared for each strain at each growth phase. The bacterial cells were sedimented by centrifugation, followed by RNA extraction, library preparation and sequencing. Ribosomal RNAs were removed with Ribo-Zero rRNA Removal Kit (Gram-Negative Bacteria). Illumina Truseq RNA libraries were prepared, with pair-end read length of 250 bp, and sequenced with Illumina Hiseq 2000 platform. The raw reads were assessed for quality, and the low-quality reads and adapters were removed. Cleaned reads were mapped to reference genomes with Bowtie (v2.3.3.1) and the default alignment settings (57). HTSeq-count was used to count the raw read counts for each gene (58).

### Comparison of gene expression profile

The pan-genome genes of the 8 *Salmonella* strains for which gene expression was profiled were retrieved from the pan-genome gene set identified from the 26 *Salmonella* strains described above. The raw reads were retrieved for each pan-genome gene from the quantification results for each growth stage, strain and replicate. The read count for a genetically missing gene was recorded as zero. Combination of the reads of pan-genome genes for all the samples generated an expression matrix, with columns and rows representing samples and pan-genome genes respectively. The gene expression levels were normalized with a model based on negative binomial distribution, and comparison was performed with edgeR between samples for the same strain under different stages or for different strains under the same stage (59). Benjamini-Hochberg corrections were performed to the genes and the significance for differential expression was preset as (1) a false discovery rate of < 0.05, and (2) fold change ≥ 3. The union set of differentially expressed genes between each pair of samples described above was analyzed, and a new expression matrix of all differential genes was prepared. The expression level for each differential gene was recorded as the count per million reads (CPM) in the corresponding sample. Primary component analysis (PCA) was performed to the differential expression matrix, followed by unsupervised clustering with the full PCA-represented features using the ‘FindClusters’ function integrated in R package Seurat v4.1.0 (60). UMAP was used to reduce the dimensions and visualize the clustering results (61). To exclude the influence exerted by genetic difference, the intersection of differentially expressed genes and core genes among the 8 strains was analyzed, and PCA and clustering were repeated with the same procedure.

### Analysis of the influence of genetic rearrangement on gene expression

Operons were inferred for each representative *Salmonella* strain based on the neighborhood relationship of the gene loci (<50 bp distance between adjacent genes) and the same direction of transcription. The 1DGR annotation results for each representative strain were further referred to, and the genes / operons and their 5’-end spanning distance away from the nearest upstream breakpoints of evolutionary blocks (noted as *distance*) were noted down. For each pair of the strains whose gene expression was profiled and compared, the shorter *distance* for the two strains was recorded for each differentially or non-differentially expressed core gene / operon. After pooling the results from all pairs of strains, comparison was performed to the distribution of *distance* between the differentially and non-differentially expressed genes / operons located in the blocks of genus origin using Mann-Whitney U tests. To analyze the influence of rearrangement scale on gene expression, the upstream nearest insertion blocks were retrieved and the block *size* was analyzed dependently on the *distance*. The *distance* was stratified and the distribution was compared between the differentially and non-differentially expressed genes / operons for the *size* of upstream blocks. The data from different growth stages were pooled, and One-way or Two-way Analysis of Covariance (ANCOVA) was performed with growth stage as a covariate. For the statistical analysis, significance level was preset as *p* < 0.05.

### Generation of *Salmonella* 3C libraries

*Salmonella* strains were inoculated from frozen stocks and grown on LB agar at 37°C overnight. A single colony was used to inoculate 5 mL of LB, and cells were grown 15h at 37°C with shaking at 200 rpm. This overnight culture was normalized to 1 OD_600_ and cells were inoculated into 100 ml of LB broth at a 1 in 1000 dilution. These cells were grown for approximately 3 h until an OD_600_ of 0.2 was reached.

To fix the proteins onto the DNA chromosome, fresh 37% formaldehyde solution was added to each culture to a final concentration of 7%; this mixture was incubated at room temperature for 30 min, then at 4°C for 30 min under gentle agitation with a magnetic stirrer. The formaldehyde was quenched by the addition of 2.5 M glycine to a final concentration of 250 mM and mixing at 4°C for 20 min. The fixed cells were pelleted by centrifugation at 4°C (3,500 x *g*, 10 min), washed with LB medium, and flash frozen in an ethanol and dry ice bath prior to storage at –80°C. Frozen cell pellets were thawed on ice for 30 min, resuspended in 600 µl 1X TE (10 mM Tris, 0.5 mM EDTA) pH 8 buffer. Lysozyme was added (5 µL of 5 mg/ml solution; approximately 2000 units), the mixture was incubated at 37°C for 20 mins, then SDS was added to a final concentration of 0.5% and the mixture was incubated at room temperature for 10 min. Each 145 µL of lysate was digested with 50 units of HpyCH4IV (Cat # R0619L, NEB, USA) for 18 h at 37°C. After digestion, the insoluble fraction of the crosslinked chromatin was sedimented by centrifugation (16,000 x *g,* 20 min) and resuspended in sterile water. The digested DNA fragments were ligated with T4 ligase (50 units/mL final concentration), in presence of NEB ligation buffer with 0.1 mg/mL of BSA and 1 mM ATP; this mixture was incubated at 16°C for 6 h. EDTA was added to stop the reaction (final concentration of 5 mM), followed by addition of proteinase K to a final concentration of 250 µg/mL and incubation at 65°C for 18 h. DNA was precipitated from solution by addition of 3M sodium acetate and isopropanol, incubation at –80°C for 1 h, and centrifugation (10,000 x g, 20 min, 4°C). This DNA pellet was allowed to air-dry, resuspended in 1x TE and RNAse A was added to a final concentration of 0.1 mg/mL, followed by incubation at 37°C for 30 min. DNA was extracted from this solution using a standard phenol: chloroform: isoamylalcohol procedure, with final resuspension in 30 µl 1x TE. The quality and quantity of each 3C library were estimated on an Agilent Bioanalyzer and by visualization on a 1% agarose gel. 5 µg of DNA for each library was resuspended in water to a final volume of 130 µL; the DNA was sonicated to break into smaller pieces. Libraries with 300-400bp inserts were prepared using NEBNext Ultra DNA library Prep kit for Illumina. Paired-end 150 bp sequencing was performed on the Illumina HiSeq platform (Novogene USA; Sacramento, CA).

### 3C sequencing data analysis

The 3C sequencing data analysis mainly followed the protocol prepared by Koszul lab for *E. coli* (25). Briefly, the raw 3C reads were filtered for the low-quality and adapter sequences, and mapped to reference genome of the corresponding *Salmonella* strain with Bowtie (v2.3.3.1) using a ‘very-sensitive’ mode (57). The scripts *fragment_attribution.py*, *library_events.py* and *Matrice_Creator.py* were run consequently to generate the intra-chromosome contact matrix file. The length of bins was set as 20-kb, 10-kb and 5-kb, respectively. Scalogram and Directional Index (DI) analysis were performed using the *multi_scale_domainogram_FILES2_dom3_3plots.py* script. The resolution was adjusted for Directional Index analysis, to find the major interval with bin size of 400kb and the CIDs with bin size of 100kb. MEME suites (62) were used to find the consensus motifs from the 0.5-kb nucleotide sequence flanking the major interval or CID borders at both sides (https://meme-suite.org/meme). E-values were calculated with MEME directly, and the significance was preset as < 0.05. For a consensus motif, MEME-ChIP (63) was used to find the homologs to the known patterns of binding sites for known transcription factors or other DNA-binding proteins (https://meme-suite.org/meme).

To find the significant long-range interaction pairs, for each target bin, exponential distributions were fit for the raw count of contacting reads for all the bins along the chromosome. With the *rate* parameter being estimated and the significance of distribution models and parameters being tested, each bin pair was further tested for the contacting significance based on the fit exponential distribution of the optimized parameter *rate*. A pair was considered with significant interaction only when the following criteria were met simultaneously: (1) the contact was significant with a Bonferroni corrected *p*-value of <0.01 using all the other genomic positions contacting with target as background; (2) except for the diagonal positions, both *x_mn_* and *x_nm_*in the raw contact read count matrix were significant, where *m* and *n* represented the order of line and column, respectively. In this study, a span of >= 50kb was considered as long range.

## Author Contributions

YW and APW conceived and supervised the project. YH and YW coordinated the project. YH, ZC, ZZ, RC and YW provided codes, models and software tools. ZZ and RC developed the website and databases. ASS and APW built the 3C libraries. YH, ZC, JZ, SL, XC, JW, YD, HB, XH, MAS and YW made the comparative genomic, gene expression and Hi-C data analysis. YH, ZC, ASS, JZ, MAS, SL, APW and YW wrote the first draft. GQZ, SLL, APW and YW revised the manuscript.

## Data and materials availability

The genome assemblies for the newly sequenced *Salmonella* strains were submitted to GenBank with accessions of CP100359 (RKS2978), CP100411 (RKS3027), CP100412 (RKS2986), CP100413 (RKS3013), CP100414 (RKS3057) and CP100415 (a plasmid of RKS3057). RNA-seq and 3C data were deposited in NCBI SRA database with accessions of SRR19426961-SRR19427008 and SRR19434835-SRR19434847 respectively. The codes and their usage for comparative genomics and 3C data analysis were accessible freely through the link: http://61.160.194.165:3080/ESG/tools or https://zenodo.org/records/10498097 or https://doi.org/10.6084/m9.figshare.24990570.

## Supplementary Materials

**Fig. S1. Phylogenomic trees of the major *Salmonella* lineages. (A)** The phylogenomic tree of major *Salmonella* lineages. The maximum likelihood tree of the genus *Salmonella* based on the amino acid sequences derived from the core gene set identified from 26 strains representing each major evolutionary branch. Bootstrapping tests were performed with 1000 replicates and the percentages of topology consistency were indicated for each node. ‘nA’, ‘nB1’, ‘nB2’, ‘nC’ and ‘nD’ were designated to represent the major nodes as shown for convenience. The phylogenetic tree based on the 16S rRNA sequences of representative *Salmonella* strains is shown in the upper left corner; this was adapted from a previous study (3). **(B)** The phylogenomic tree of representative *S. enterica subsp. enterica* serovars. A procedure similar to **(A)** was used for the phylogenomic building and visualization. *S. enterica subsp. indica* strain RKS3057 was used as an outgroup to determine the root of the serovars. The strains were clustered into 4 major clusters (Clusters 1∼3 and the Hillingdon group), while Cluster 1 were further classified into 11 sub-clusters, namely 1a through 1k.

**Fig. S2. Characterization of genetic rearrangements within the *Salmonella* genus. (A)** The distribution of spontaneous rearrangements (one-time) and hot spots (2-14 times) of rearrangement were calculated. **(B)** The functional distribution is shown for the rearranged operons from the *Salmonella* ancient orthologous chromosomes.

**Fig. S3. Evolution of the *lpf* operon in *Salmonella*. (A)** Conserved gene order and synteny of *lpf* locus between *S. enterica subsp. enterica* LT2 and *S. bongori* RKS3044. **(B)** – **(C)** Collinearity of *lpf* locus and the flanking 20-kb nucleotide sequences at both upstream and downstream between LT2 and RKS3044 **(B)** or between the ancient orthologous genomes of *S. enterica subsp. enterica* and *S. bongori* **(C)**. The *lpf* locus was indicated within the red boxes. The collinearity analysis was performed using PipMaker (64).

**Fig. S4. Evolution of *cdtABC* locus in *Salmonella*. (A)** Conserved gene order and synteny *cdtABC* locus between *S. arizonae* RKS2980 and *S. diarizonae* 11_01855. **(B)** – **(C)** Collinearity of *cdtABC* locus and the flanking 20-kb nucleotide sequences at both upstream and downstream between RKS2980 and *S. diarizonae* 11_01855 **(B)** or between the ancient orthologous genomes of the *Salmonella* genus and *S. enterica* **(C)**.

**Fig. S5. The collinearity analysis of typhoid toxin and flanking loci in different *Salmonella* strains**. For each strain, the typhoid toxin (TT) encoding and 5-kb flanking nucleotide sequences in both sides were retrieved and aligned with others using PipMaker (64). The *TT* locus containing *artA*, *artB* and adjacent genes, is not as conserved for gene composition, gene order and genomic loci among the positive strains from *S. bongori*, *S. enterica subsp. arizonae* and *S. enterica subsp. diarizonae* **(A-C)**. However, the *TT* loci within *S. enterica subsp. enterica* serovars, such as Paratyphi A and Javiana, showed good collinearity between each other, and with *S. bongori* strains **(D-E)**. The loci within the *S. enterica subsp. enterica* serovars and *S. bongori* strains were also collinear with serovar Typhi **(F)**, but the *TT* loci in Typhi was located at the border of a large-scale genomic inversion so it only shows the appearance of single-end collinearity. Given the frequent hot-spot rearrangement loci within *Salmonella* genomes, the sporadic detection of *TT* in other serovars (65), and different flanking sequences, presumably of phage origin, the *TT* loci were most likely not present in the *Salmonella* genus ancient genome but rather acquired by the subsequent phylogenies independently.

**Fig. S6. Differential gene expression profiles among *Salmonella* strains.** The numbers of all differentially expressed genes **(A)** or differentially expressed genes that are part of the *Salmonella* core genome **(B)** are listed between different strains at the same growth phase (i.e., exp, exponential (red background); stat, stationary (green background)), or between the same strain at different growth phases (i.e., yellow background).

**Fig. S7. The consistency and correlation of 3C contacts between replicates of the *Salmonella* strains**. **(A)** Consistency of 3C contact maps between biological replicate samples (shown top to bottom for each strain). The bin size was set as 5 kb. **(B)** Correlation of the 3C contacts between biological replicates for each strain was calculated at 5 kb, 10 kb and 20 kb bin sizes.

**Fig. S8. Boundary variation of *Salmonella ter* interaction interval**. **(A)** *Salmonella* interaction intervals analyzed according to the 400-kb DI results. **(B)** 400-kb DI analysis of chromosomal contact map of MG1655. The boundaries of interaction intervals analyzed according to the 400-kb DI results were indicated above the DI diagrams, and the homologous pairs between MG1655 and *Salmonella* strains were connected with lines. **(C)** The position of sequence homologs of MG1655 *ter* interaction interval boundaries in the chromosomes of *Salmonella* strains. The positions were indicated with red lines. **(D)** Diagrams illustrate the comparative genomic analysis between MG1655 and each individual *Salmonella* strain. Pink blocks represent inverted segments, whereas the black and purple blocks represent homologous regions between the strains compared.

**Fig. S9. Consensus motifs identified from the conserved interaction interval boundaries of *Salmonella* and *E. coli* MG1655.** The motifs were screened with MEME (63) (https://meme-suite.org/meme), and the E-values were indicated.

**Fig. S10. The distribution of *ter* interaction interval in the ancient orthologous chromosome of *Salmonella* genus.** The *ter* interaction interval was shown in orange in the outmost circle. The hotspot and spontaneous rearrangement loci were shown in red and blue respectively in the middle circle, and the GC-skew curves were shown in the inner circle.

**Fig. S11. Sequence homology of the *Salmonella* CID boundaries. (A)** Sequence homology of the CID boundaries between the four *Salmonella* strains analyzed. The conserved pairs were connected with solid lines. **(B)** Sequence homology in the RKS3044 chromosome for the CID boundaries in other *Salmonella* strains but missed in RKS3044. The homologous pairs were connected with solid lines.

**Fig. S12. Consensus motifs identified from the boundaries of chromosome interaction domains and/or interaction intervals of *Salmonella* and *E. coli* MG1655.** The motifs were screened with MEME (63) (https://meme-suite.org/meme), and the E-values were indicated.

**Fig. S13. Comparison of *ter* interaction interval sequences between RKS3057 and other *Salmonella* strains**. MD, Interaction interval. **(A)** The SPI-2 T3SS is located around the upper boundary of *ter* interaction interval of RKS3057. In RKS3044, the SPI-2 T3SS is missing but a T6SS is encoded in the same locus. The fragment length of the T3SS in RKS3057 and the T6SS in RKS3044 is similar and the flanking sequences appear conserved. However, the SPI-2 T3SS in RKS3057 is contained in the *ter* interaction interval while the T6SS from RKS3044 is outside, possibly due to different long-range interactions of the CIDs with nearby ones. **(B)** The non-conservation of the upstream boundaries of the CIDs where SPI-2 T3SSs are located between RKS3057 and RKS3027 could be related with the two large insertions/deletions (∼14 kb together) within the regions flanking the conserved boundary sequences. **(C)** RKS2980 shows a ∼80kb inversion at the downstream of SPI-2 T3SS with RKS3057, and correspondingly, the composition and boundaries of CIDs within the *ter* interaction interval generally show large differences. Mauve v2.4.0 (55) was used to perform the comparison and to generate the diagrams.

**Fig. S14. Intra-chromosomal long-range interaction profiles of repeated hubs.** The interactions shown between loci occurred over a span of >50kb. The interaction profiles are shown for RKS3044 **(A)**, RKS2980 **(B)**, RKS3027 **(C)** and RKS3057 **(D)**. Each line connects the long-range interacting partners. One partner from an interacting pair represented repeated hub, which was shown in colored line within the outer circle and the encoded genes were indicated. The reference chromosome sequence of each *Salmonella* strain was adjusted to match the annotation start site and strand referred to in *E. coli* MG1655. SPI-2 T3SS’ and *sipB/sseJ’* in RKS3044 represent the genomic coordinates where the SPI-2 T3SS and *sipB/sseJ* are located in *S. enterica* species respectively.

## Datasets S1 to S7

(The datasets can be accessed at https://zenodo.org/records/10498097)

**Dataset S1. Salmonella genomes newly sequenced and used in this study.**

**Dataset S2. The core gene set of 26 representative *Salmonella* strains used for manual AOC construction.**

**Dataset S3. Genes and their evolutionary routes inferred from the evolutionary trajectory of *Salmonella* ancient orthologous chromosomes.**

**Dataset S4. Normalized expression levels of genes for the pan gene sets of *Salmonella* or common gene sets among the eight representative strains.**

**Dataset S5. Evolutionary origins of differentially expressed genes.**

**Dataset S6. Consensus sequence motifs identified from the flanking (+/− 500bp) regions of the major interaction intervals or CID boundaries.**

**Dataset S7. Presence of a large inversion spanning the *ter* major interval between *E. coli* lineages and *Salmonella*.**

